# *Culex pipiens* L. and *Culex restuans* egg rafts harbor diverse bacterial communities compared to their midgut tissues

**DOI:** 10.1101/2020.05.23.112128

**Authors:** Elijah O. Juma, Chang-Hyun Kim, Christopher Dunlap, Brian F. Allan, Chris M. Stone

**Author notes:** Corresponding Author: 505 S. Goodwin Ave, Urbana, IL 61801, USA.

## Abstract

**Background:** The bacterial communities associated with mosquito eggs are an essential component of the mosquito microbiota, yet there are few studies characterizing and comparing the microbiota of mosquito eggs to other host tissues.

**Methods:** We sampled gravid female *Culex pipiens* and *Culex restuans* from the field, allowed them to oviposit in the laboratory, and characterized the microbiota associated with their egg rafts and midguts for comparison through MiSeq sequencing of the 16S rRNA gene.

**Results:** Bacterial richness was higher in egg rafts than in midguts for both species, and higher in *Cx pipiens* L. than *Cx. restuans*. The midgut samples of *Cx. pipiens* and *Cx. restuans* were dominated by *Providencia. Culex pipiens* L. and *Cx. restuans* egg rafts samples were dominated by *Ralstonia* and *Novosphingobium*, respectively. NMDS ordination based on Bray-Curtis distance matrix revealed that egg raft samples, or midgut tissues harbored similar bacterial communities regardless of the mosquito species. Within each mosquito species, there were distinct clustering of bacterial communities between egg raft and midgut tissues.

**Conclusion:** These findings expand the list of described bacterial communities associated with *Cx. pipiens* L. and *Cx. restuan*s and the additional characterization of the egg raft bacterial communities facilitates comparative analysis of mosquito host tissues, providing a basis for future studies seeking to understand any functional role of microbiota in mosquito biology.

## Introduction

Studies applying high throughput, culture-independent sequencing of the bacterial 16S rRNA gene have advanced understanding of the association between mosquitoes and their bacterial communities [1–4]. The bulk of studies characterizing mosquito-associated bacterial communities have focused on the mosquito gut. However, other mosquito organs or tissues, including the ovaries, the male reproductive system, the salivary glands, and eggs, are also known to harbor bacterial communities that may play essential roles in mosquito biology [5–11]. The research focus on the mosquito gut, especially in the adult stage, is underpinned by the understanding that the mosquito midgut environment typically is the first barrier that mosquito-borne pathogens must overcome to develop successfully within the mosquito host and be transmitted to the next susceptible host [12–15].

Mosquito egg bacterial communities likely also play important roles in mosquito ecology. Studies with *Aedes aegypti* and *Ae. triseriatus* have shown that bacterial density on the egg surface mediates egg hatch rates and time-to-hatch, while mosquito eggs subjected to heavy larval grazing in high larval density habitats can exhibit delayed time-to-hatch [16–18]. The presence of aerobic microorganisms on the egg surface and in the larval environment has been associated with reduced oxygen tension, providing the stimulus for mosquito egg hatching [19,20]. To what extent the bacterial communities on the egg surface drive the hatching effect relative to the microbes in the water column remains unclear.

Previous studies characterizing and comparing the bacterial communities of mosquito eggs with those of their other host tissues (e.g., midguts) have been conducted with well-known Afro-tropical or Asian vectors, including *Anopheles* or *Aedes* species, but not important North American vector species such as *Culex pipiens* L. or *Culex restuans* [5–7]. *Culex pipiens* L. is an introduced European species that arrived in North America in the early 16^th^ century through trade and has been naturalized in the United States north of 39 ° latitude [21–23]. It serves as both an amplifying and bridge vector for West Nile virus (WNV) and St. Louis encephalitis due to its preference for feeding on birds [24]. *Culex restuans*, native to North America, is also an important vector of WNV and distributed in the northeast and Great Lakes regions of the United States [22,24]. These container-dwelling mosquitoes are common in urban, residential neighborhoods and woodlots [25–27]. Comparative studies with *Cx. pipiens* and *Cx. restuans* are necessary to expand the known library of bacterial communities associated with mosquito host tissues and to facilitate further studies on the role of these bacterial communities in mosquito vector biology and ecology and potentially benefit mosquito-borne disease control.

We used Illumina MiSeq sequencing of the V3-V4 hypervariable regions of the 16S rRNA gene to characterize the bacterial communities associated with egg rafts and midguts of *Cx. pipiens* L. and *Cx. restuans*, to gain insights into bacterial community structure and diversity, and to observe how they compare between the two host tissues. We tested the hypothesis that, within each mosquito species, egg raft and the midgut samples will harbor distinct bacterial communities given that the external egg surface and the mosquito midgut represent physiologically distinct environments, and thus are likely to support distinct bacterial communities. We also hypothesized that similar tissues across species (e.g. eggs or midguts for both *Cx. pipiens* and *Cx. restuans*) will harbor similar bacterial communities because similar tissues represent similar physiological environments. This study expands upon understanding of the bacterial communities associated with mosquito host tissues other than the mosquito gut and provides a basis for further studies focused on the role of mosquito egg-associated bacterial communities in mosquito biology and mosquito-borne disease control.

## Materials and methods

### Sampling and laboratory sample preparation

Gravid traps for sampling of gravid *Culex* spp. mosquitoes were established in three woodland areas namely, Brownfield Woods (40° 8’ 46.0716’’ N, 88° 9’ 57.0852’’ W), South Farms (40° 5’ 18.7692’’ N, 88° 13’ 0.4188’’ W), and Trelease Woods (40° 7’ 45.5412’’ N, 88° 8’ 28.2696’’ W), and two residential neighborhoods with permission from property owners (40° 4’ 57.0324’’ N, 88° 15’ 25.7652’’ W; 40° 5’ 25.8828’’ N, 88° 15’ 36.4212’’ W) in Champaign County, Illinois. At each sampling site, two CDC gravid traps baited with 3.8 L each of grass infusion [28] were deployed beginning on June 18, 2018 and sampling was conducted three times weekly up to July 20, 2018. Traps were placed in the evening just before dusk and the collection bags collected approximately 12 hours later the next morning [29]. Individual gravid female *Culex* species mosquitoes were transferred separately to individual 270 mL paper cups to facilitate oviposition; each consisted of a 30 mL inner plastic oviposition cup filled to half capacity with distilled water. The mosquitoes were maintained in a walk-in environmental chamber at 27°C±1 and ∼75±5% relative humidity with a 16:8 (L:D) photoperiod. The paper cups were provisioned with cotton balls soaked in distilled water to provide additional humidity and a source of water for the gravid females. Monitoring for egg rafts was conducted every 12 hours. Following oviposition, the egg raft and the parous female were separately preserved at −80°C for future bacterial DNA extraction.

### Dissections & DNA extraction

Egg rafts and mosquito samples were thawed and adult mosquito samples surface sterilized in 70% ethanol for 5 minutes, transferred to 3% bleach solution for 3 minutes, transferred again to 70% ethanol for 5 minutes, and then rinsed 3 times in sterile water and 4 times in Dulbecco’s Phosphate-Buffered Salines (DPBS) (ThermoFisher Scientific, Waltham, MA) [4]. Each sample was dissected in a small drop of sterile DPBS using a stereo-dissecting microscope and the midguts were transferred to PowerSoil bead tubes. Similarly, egg rafts were individually transferred to PowerSoil bead tubes. Due to their hydrophobicity and ease of disintegration during handling, egg rafts were processed for bacterial DNA without surface sterilization. Samples were homogenized using Retsch MM 300 TissueLyser (Retsch, Haan, Germany) and genomic DNA was extracted using MoBio PowerSoil DNA Isolation Kit (MoBio Laboratories, Inc., CA) according to the manufacturer’s instructions. DNA was quantified using the Nanodrop 1000 (ThermoFisher Scientific, Pittsburgh, PA).

Sequencing was performed at the National Center for Agricultural Utilization Research, Peoria, IL. The V3-V4 hypervariable region of bacterial 16S rRNA gene was PCR-amplified using previously published universal primers 341f and 806r [30,31]. The V3-V4 hypervariable region has been shown to have higher sensitivity in bacterial phylogenetic analysis compared to the rest of the hypervariable regions of the 16S rRNA gene [1]. The following primer set specific for the V3-V4 region of the 16S rRNA gene was used: Forward 5’CCTACGGGNGGCWGCAG; Reverse 5’GACTACHVGGGTATCTAATCC. The primers were incorporated into fusion primers for dual indexing and incorporation of adapters prior to genome sequencing using Illumina MiSeq (Illumina Inc., San Diego, CA) [32]. The V3-V4 hypervariable region of the bacterial 16S rRNA gene was PCR-amplified using the following primer set: Forward 5’CCTACGGGNGGCWGCAG; Reverse 5’GACTACHVGGGTATCTAATCC. PCR was done in 25 µL reactions containing 12.5 µL of 2x KAPA HiFi HotStart ReadyMix, 5 µL of 1 µM each of the forward and reverse primers, and 2.5 µL of template genomic DNA. PCR conditions were 95 °C for 3 min; 25 cycles of: 95 °C for 30s, 55 °C for 30s, 72 °C for 30s; 72 °C for 5 mins; hold at 4 °C. PCR amplicons were cleaned using AMPure XP beads to remove free primers and primer-dimer species. A second PCR was conducted using the Nextera XT Index Kit (Illumina, San Diego, CA) to attach dual indices and Illumina sequencing adapters. Index PCR was conducted in 45 µL reactions containing 25 µL of 2x KAPA HiFi HotStart ReadyMix, 5 µL each of index 1 and index 2 combinations, and 10 µL of PCR grade water. Thermocycling conditions were 95 °C for 3 min; 8 cycles of 95 °C for 30s, 55 °C for 30s, 72 °C for 30s; 72 °C for 5 mins; hold at 4 °C. A negative control sample made up of DNA extracted from molecular biology grade water was sequenced with the same protocol to allow detection of the contamination background. PCR amplicons were cleaned and normalized using a SequalPrep normalization plate (Thermofisher Scientific, Waltham, MA). The pooled library was mixed with Phix control spike-in of 5% as a sequencing control. The samples were sequenced on Illumina MiSeq system with a MiSeq V3 2 × 300 bp sequencing kit. The demultiplexed reads were quality-trimmed to Q30 using CLC genomics workbench v12.0 (Qiagen Inc., Valencia, CA). Read pairing, fixed-length trimming and OTU clustering were done using CLC Bio Microbial Genomics module (Qiagen Inc., Valencia, CA) utilizing the reference sequences from the Greengenes ribosomal RNA gene database [33]. The operational taxonomic unit (OTU) assignment was done at 97% sequence similarity, which is considered adequate for bacterial identification to the genus level [34].

### Species identification

A duplex real-time TaqMan PCR assay [35] was used for molecular identification of *Cx. Pipiens* L. and *Cx. restuans* using primers and probes targeting the *acetylcholinesterase* gene (*Ace2*) adopted from [35]. *Culex pipiens* L. primers and probes consisted of: CxPip-F1 GGTGGAAACGCATGACCAGATA; CxPip-R1 TGCAATAAAGAGGTGGCCACG; and probe FAM/AGCCACGAACAACTAAATCATCACAAGCACAGC/3BHQ. *Culex restuans* primers and probes were made up of: CxRest-F1 ATCGGTCTGGCTTCCTTTCAGAT; CxRest-R1 TTAGTCAAGTTAACTGGCCTACATCCTA, and the probe JOE/AGCAAACTGGCCGTCGTCCACCGATATAAAT/3BHQ_1. The target DNA used as a template was taken from DNA samples extracted from midgut samples for bacterial DNA analysis. Separate DNA samples were extracted from *Cx. pipiens* and *Cx. restuans* adults initially identified from larval stages and used as positive controls. Additionally, a reaction mixture consisting of *Ae. albopictus* DNA template minus reverse transcriptase was used as a negative control to rule out any chance of contamination in the PCR reaction. Each PCR sample was assayed in 25 µL reaction mixture consisting of 5 µL of the target DNA, 12.5 µL SensiFAST(tm) Probe Hi-ROX Kit master mix (Bioline, Tauton, MA), 1.25 µL each of *Cx. pipiens* forward and reverse primer; 0.625 µL *Cx. pipiens* probe (Fam); 1.25 µL each of *Cx. restuans* forward and reverse primer; 0.625 µL *Cx. restuans* probe (Fam) and 1.25 µL of nuclease-free water. Thermocycling was performed on an ABI 7300 HT sequence detection system (Applied Biosystems, Foster City, CA) using the following reaction conditions: 95°C for 5 min followed by 40 cycles of 95°C for 30 s, 58°C for 30 s, and 72°C for 60 s [35].

### Statistical analysis

All analyses were conducted using R version 3.6.1 [36] within the Rstudio environment version 1.2.1335 [37] and PAST version 3.15 [38]. OTUs accounting for <0.005% of the total number of sequences were removed prior to analysis to eliminate spurious OTUs [39]. The OTU sequence numbers varied markedly within samples (mean ± SE = 7,621.28±547.17 per sample). Bacterial sequences were rarefied to an even depth of 1007 reads per sample to standardize the sampling coverage [40,41]. From an initial sample size of 188 samples, 44 samples had <1007 reads and were excluded from further analysis. To estimate sample coverage, rarefaction curves were fitted on the unrarefied data using the “phyloseq” package version 1.24.0 in R [40–42]. Alpha diversity metrics, including Shannon diversity index, observed species, and chao1, were generated in QIIME 2 [43]. The means and 95% confidence intervals were calculated in R to test for significant differences in the alpha diversity indices between treatments. The Kruskal-Wallis test was used to test for differences in means between egg raft and midgut samples and the Wilcoxon rank sum test with Bonferroni correction was performed pairwise to separate significant treatments. Beta-diversity measures were estimated using the Bray-Curtis dissimilarity index using the “phyloseq” package and non-metric multidimensional scaling (NMDS) ordination plots were generated to visualize the results. The non-parametric Analysis of Similarity (ANOSIM) test with Bonferroni-adjustments was performed in PAST version 3.26 [38] to determine degree of dissimilarity in bacterial composition between treatment groups. Similarity percentage (SIMPER) analysis was performed in PAST to identify the bacterial species characterizing each treatment group. Venn diagrams were generated using the R package “limma” [44] version 3.40.2 to visualize OTUs that were shared between egg rafts and midgut samples of *Culex pipiens* L. and *Culex restuans*. Using the function chisq.test from the R package “RVAidemMemoire” version 0.9-74, differences in bacterial OTU per sample were tested and pairwise multiple comparison test with Bonferroni-correction was applied.

## Results

### Sequence processing and alpha diversity analysis

Sequencing of the V3-V4 regions of the 16S rRNA gene from 188 samples (66 *Cx. pipiens* egg rafts; 66 *Cx. pipiens* midguts; 28 *Cx. restuans* egg rafts; 28 *Cx. restuans* midguts) generated 1,432,800 raw sequences (mean ± SE = 7,621.28±547.17 per sample). After quality-filtering to remove chimeric sequences, other non-bacterial sequences, and bacterial OTUs constituting <0.005% of the total sequences and rarefying the reads to an even depth of 1,007 sequences per sample to standardize sampling effort, a total of 144 samples were retained (59 *Cx. pipiens* egg rafts; 39 *Cx. pipiens* midguts; 27 *Cx. restuans* egg rafts; 19 *Cx. restuans* midguts). This sample size constituted a total of 1,422,059 sequences (mean ± SE = 9,875.41±598.19 per sample) clustered into 153 bacterial OTUs and assigned taxonomic identity at 97% sequence similarity.

Rarefaction analysis of the bacterial OTU samples revealed that the sequencing depth coverage sufficiently recovered most of the bacterial OTUs. Chao1 estimator revealed that up to 86.3%±0.07 (mean±SE) of the bacterial OTUs were recovered. The highest bacterial OTU richness was reported in *Cx. pipiens* eggs, while *Cx. restuans* midgut had the lowest bacterial OTU richness (**Additional file 1: Fig. S1**). *Culex pipiens* egg raft samples had significantly higher observed and expected (Chao1) bacterial OTU richness compared to *Cx. pipiens* midgut samples, or *Cx. restuans* egg raft and midgut samples. *Culex restuans* egg samples had significantly higher observed and expected (Chao1) bacterial OTU richness compared to *Cx. restuans* midgut samples (Observed OTUs: Kruskal-Wallis chi-squared = 92.72, df = 3, *p* < 0.0001; Chao1: Kruskal-Wallis chi-squared = 89.02, df = 3, *p* < 0.0001; Shannon index: Kruskal-Wallis chi-squared = 85.82, df = 3, *p* < 0.0001) (**Table 1**).

**Table 1.** Bacterial OTU richness and diversity (mean ±SE) in midgut and egg samples of *Cx. restuans* and *Cx. pipiens* mosquito species; n1 – sample size used in sequencing; n2 sample size retained after quality checks of the sequenced samples and used in the analysis

| Sample type | n1 | n2 | Observed | Chao1 | Shannon | OTU<sub>obs</sub>/OTU<sub>pred</sub> |
| --- | --- | --- | --- | --- | --- | --- |
| *Culex pipiens* L. midgut | 66 | 59 | 17.5±1 | 20.7±1 | 1.1±0.1 | 0.86±0.02 |
| *Culex pipiens* L. eggs | 66 | 39 | 48.6±2 | 56.9±2 | 2.9±0.1 | 0.86±0.01 |
| *Culex restuans* midgut | 28 | 27 | 15.5±2 | 19.1±2 | 0.8±0.2 | 0.88±0.02 |
| *Culex restuans* eggs | 28 | 19 | 44.3±3 | 50.9±3 | 2.7±0.1 | 0.87±0.3 |

### Taxonomic classification and bacterial composition

The 153 bacterial OTUs were classified into 7 phyla, 13 classes, 27 orders, 40 families, and 54 genera. The most dominant phyla were Proteobacteria (85.5%), consisting of Alphaproteobacteria (34.0%), betaproteobacteria (18.4%), gammaproteobacteria (33.0%), and deltaproteobacteria (0.01%). Other phyla included Spirochaetes (9.3%), Bacteroidetes (3.3%), and Firmicutes (1.1%), and the rest were <1% cumulatively (**Fig. 1A)**. Alpha- and betaproteobacteria were dominant in egg raft samples of either species, while gammaproteobacteria was dominant in the midgut samples of both species. Spirochaetes was abundant in *Cx. restuans* midgut samples (**Fig. 1A)**. The top five most abundant families accounted for 68.7% of all sequences. They included enterobacteriaceae (25.8%), sphingomonadaceae (15.9%), oxalobacteraceae (10.6%), borreliaceae (9.3%), and rickettsiaceae (7.1%). Enterobacteriaceae was dominant in the midgut samples of both species, sphingomonadaceae in *Cx. restuans* egg samples, oxalobacteraceae in *Cx. pipiens* egg samples, and borreliaceae in *Cx. restuans* midgut samples (**Fig. 1B)**. At the genus level, the top five most abundant OTUs accounted for 58.1% of all sequences. They included *Providencia* (17.8%), *Novosphingobium* (13.6%), *Ralstonia* (10.3%), *Spironema* (9.3%), and *Wolbachia* (7.1%) (**Fig. 1C)**. *Providencia* was the dominant taxon in the midgut samples of both species, *Novosphingobium* in *Cx. restuans* egg samples, *Ralstonia* in *Cx. pipiens* egg samples, and *Spironema* in *Cx. restuans* (**Fig. 1C)**. Overall, 64 (41.8%) bacterial OTUs were shared between all sample type combinations of mosquito species and host tissue (**Additional file 2: Fig. S2A**). Seventy-two bacterial OTUs (61.5%) were shared between *Cx. pipiens* midgut samples and *Cx. restuans* midgut samples and 128 of the bacterial OTUs (91%) were shared between the egg samples of the two mosquito species. Ninety-three (62.8%) bacterial OTUs were shared between egg and midgut samples of *Cx. pipiens*, while 73 bacterial OTUs (51.4%) were shared between egg and midgut samples of *Cx. restuans* mosquitoes. Overall, there was higher bacterial OTU richness and diversity in egg rafts compared to midgut samples for both *Cx. pipiens* and *Cx. restuans*. One hundred and five bacterial OTUs were detected in *Cx. pipiens* midgut samples (CXP.MG) compared to 138 in *Cx. pipiens* egg raft samples (CXP.EG), and 83 and 132 in *Cx. restuans* midgut samples (CXR.MG) and *Cx. restuans* egg raft samples (CXR.EG), respectively. The differences in OTUs detected per sample were statistically significant (Chi-squared = 17.0, df = 3, *p* < 0.001). Multiple pairwise comparison with Bonferroni corrections revealed two statistically different sample groups (CXP.EG vs. CXR.MG, *p*=0.001; CXR.MG vs. CXR.EG, *p*=0.005). Non-metric multidimensional scaling (NMDS) using Bray-Curtis distance matrix revealed bacterial communities clustered by host tissue and are supported by results of ANOSIM pairwise comparisons: (CXP.EG vs. CXR.EG; ANOSIM: *R*=0.19, *p*<0.001), (CXP.MG vs. CXR.MG (ANOSIM: *R*=0.30, *p*<0.001) (**Fig. 2, Table 2**). *Culex restuans* egg raft and midgut samples formed distinct clusters on the NMDS plot indicating distinct bacterial community composition (CXR.EG vs. CXR.MG; ANOSIM: *R*=0.70, *p*<0.001), whereas there was moderate overlap in *Cx. pipiens* egg raft and midgut bacterial communities, but still formed distinct clusters (CXP.EG vs. CXP.MG; ANOSIM: *R*=0.51, *p*<0.001) (**Fig. 3, Table 2**). SIMPER analysis identified 9 bacterial OTUs that were responsible for 70% of the observed differences between groups, with *Providencia* (19.14%), *Ralstonia* (10.57%), *Novosphingobium* (10.34%), and *Spironema* (8.39%) constituting the largest variation (**Additional file 3: Table S1**).

**Fig. 1.**
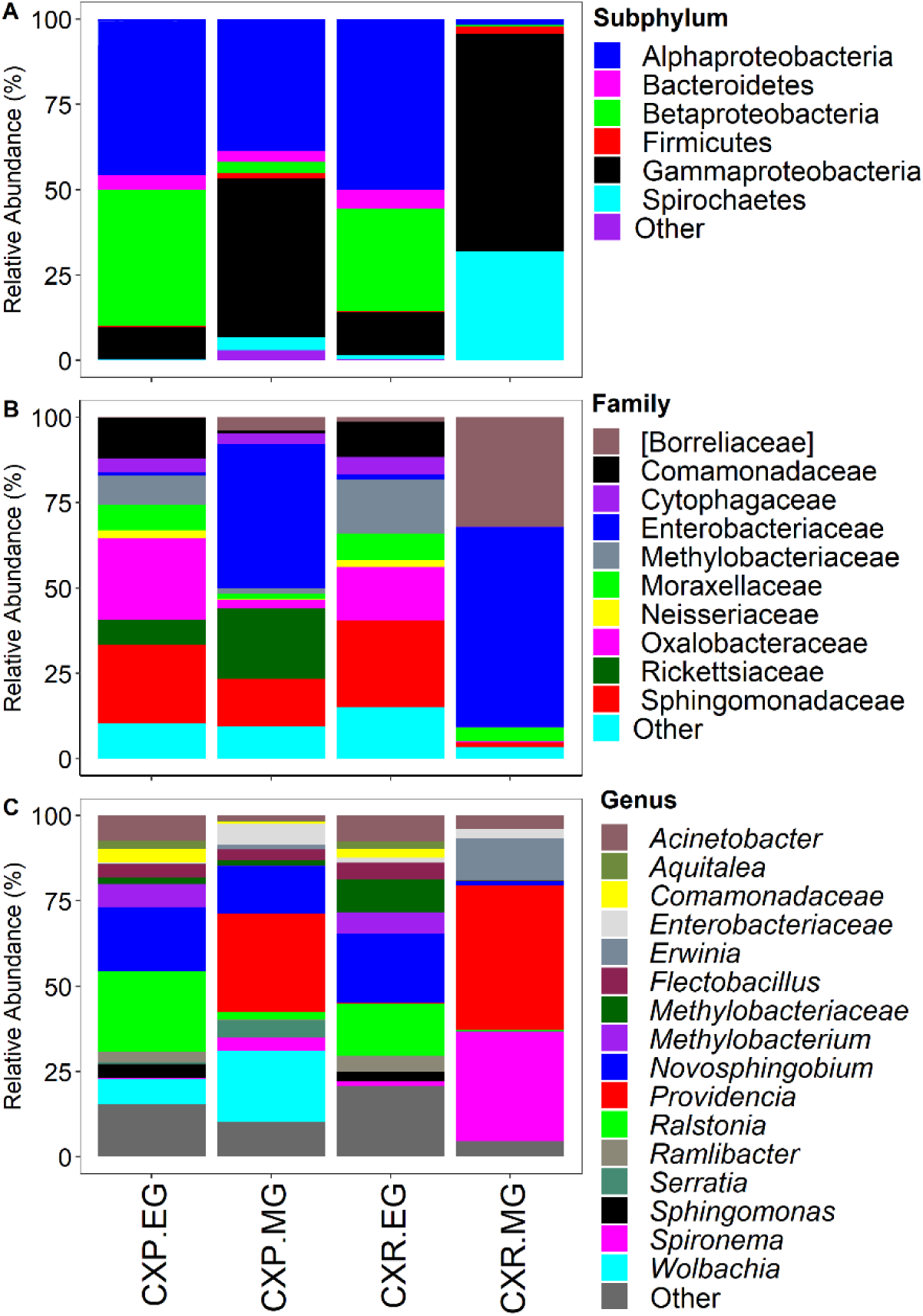
Relative abundance of bacterial communities in samples of *Cx. pipiens* and *Cx. restuans* midgut and egg samples. Taxa with sequence abundance <1% of total sequences were pooled together as “Other” in all the taxonomic ranks. CXP.EG – *Cx. pipiens* egg raft samples; CXP.MG – *Cx. pipiens* midgut samples; CXR.EG – *Cx. restuans* egg raft samples; CXR.MG – *Cx. restuans* midgut samples

**Table 2.**
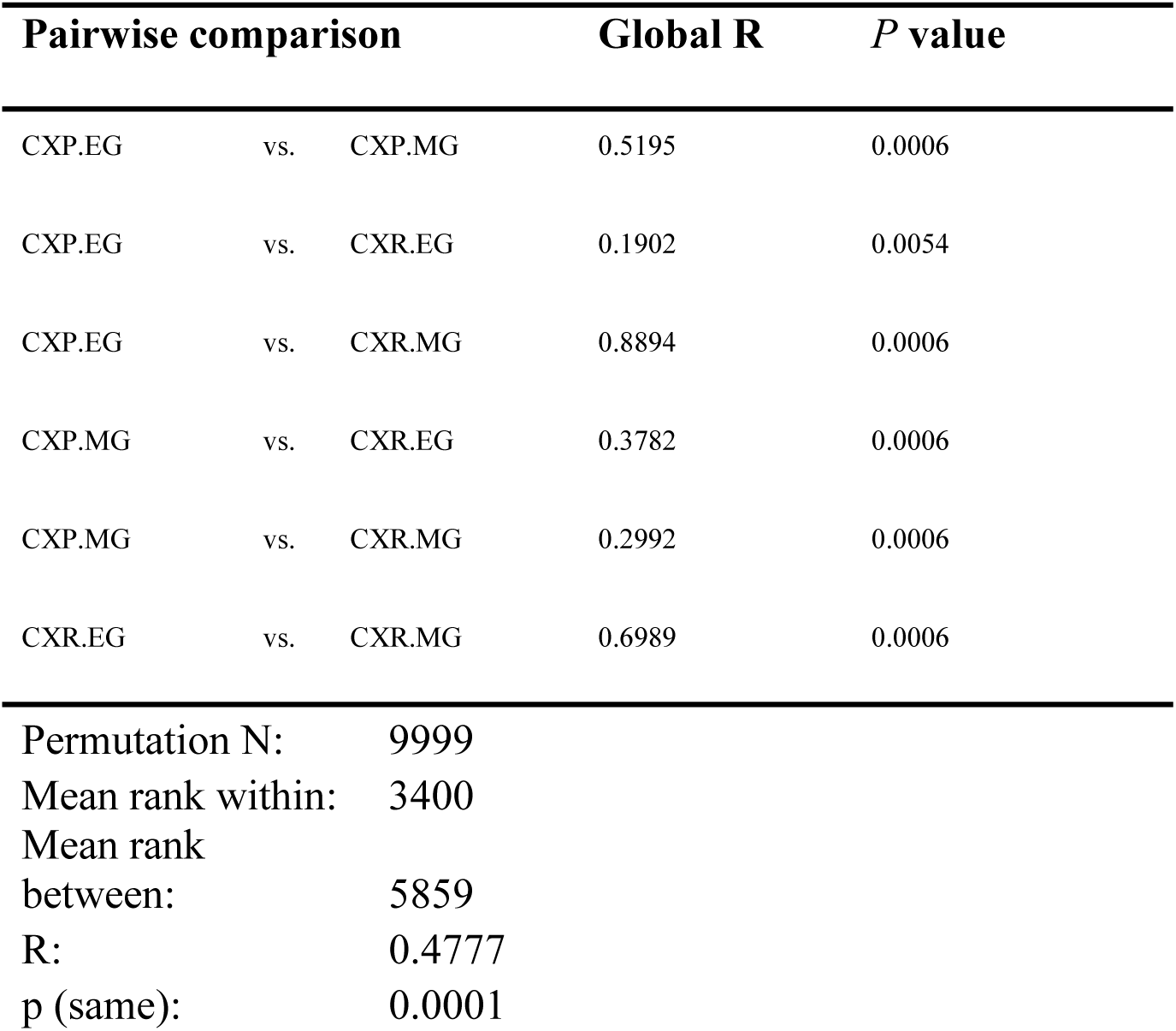
Pairwise ANOSIM comparisons by mosquito species and life stage. The significance values are Bonferroni-corrected for multiple comparisons CXP – *Cx. pipiens*; CXR – *Cx. restuans*; EG – egg samples; MG – midgut samples

**Fig. 2.**
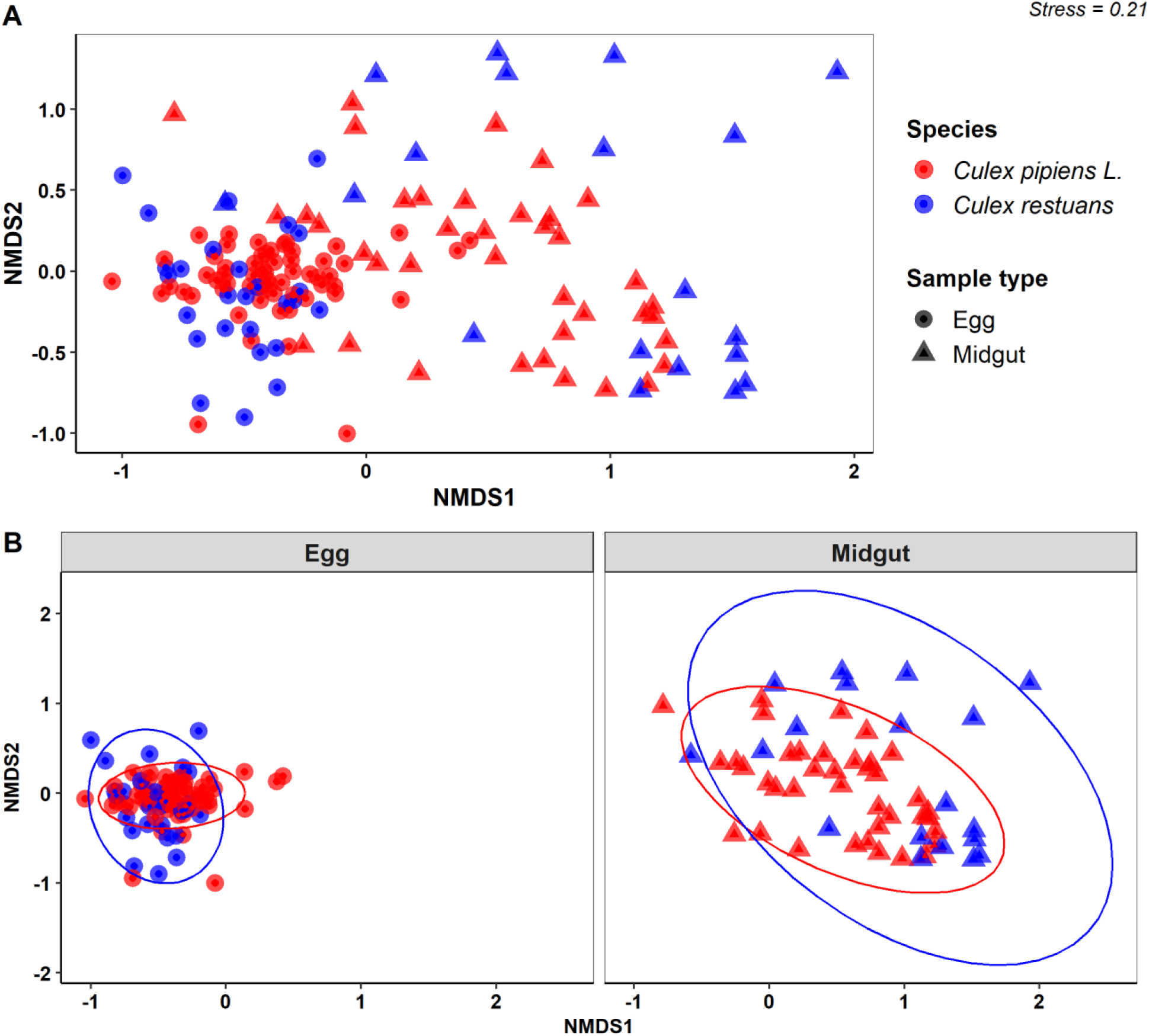
NMDS based on Bray-Curtis distance matrix of bacterial communities from *Cx. pipiens* and *Cx. restuans* egg and midgut samples. **A:** Bacterial communities from *Cx. pipiens* and *Cx. restuans* samples presented together; **B:** Bacterial communities from *Cx. pipiens* and *Cx. restuans* samples partitioned by life stage sampled

**Fig. 3.**
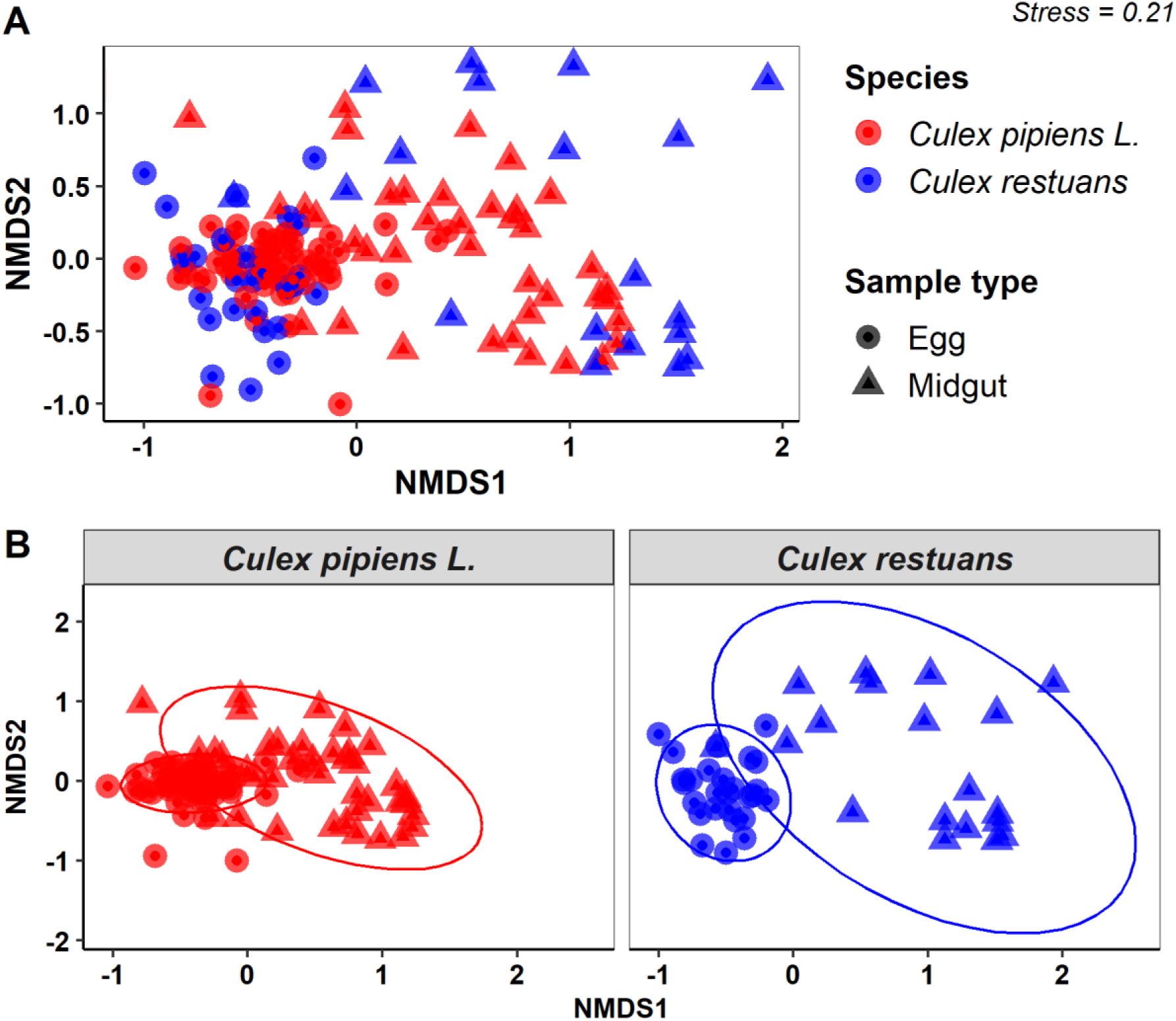
NMDS based on Bray-Curtis distance matrix of bacterial communities from *Cx. pipiens* and *Cx. restuans* egg and midgut samples. **A**: Bacterial communities from *Cx. pipiens* and *Cx. restuans* samples presented together; **B**: Bacterial communities from egg and midgut samples partitioned by the mosquito species

*Providencia* was the most dominant bacterial OTU in midgut samples of both *Cx. pipiens* and *Cx. restuans. Ralstonia* was the dominant bacterial OTU in *Cx. pipiens* egg raft samples but was also present in high proportions in *Cx. restuans* egg raft samples. *Novosphingobium* was dominant in egg raft samples from both species. *Spironema* was the dominant bacterial taxon in *Cx. restuans* midgut samples (**Fig. 1C**).

## Discussion

In this study, we characterized the composition and diversity of the bacterial communities associated with egg rafts and midgut samples of *Cx. pipiens* and *Cx. restuans*. The egg raft samples in both species were more diverse compared to midgut samples, with *Cx. restuans* midgut samples supporting the lowest bacterial diversity. Bacterial communities clustered by mosquito host tissue, such that the egg rafts from *Cx. pipiens* and *Cx. restuans* shared substantially similar bacterial communities and so did their midgut samples. However, both species had significantly different bacterial communities between their egg raft and midgut tissues.

The bacterial communities associated with mosquito eggs are mostly localized on the external surfaces of the eggs. Previous studies have not been able to isolate bacterial communities from within the egg cytoplasm [4,6,7,17,45]. The egg raft samples from both species in our study had highly overlapping bacterial communities with up to 91% of the bacterial OTUs shared between them. We presume, based on existing evidence, that these bacterial communities were mostly localized on the exterior of the egg rafts, thus representing the natural bacterial communities supported by the exterior egg raft surface environment. In this study, female mosquito oviposition took place in deionized water, which is deficient in microorganisms [46], and the egg rafts were preserved at −80 °C within hours of oviposition.

These measures limit the possibility that any significant level of bacterial colonization of the egg rafts may have occurred immediately following oviposition. We suspect that most of the bacterial communities colonizing the egg rafts were inherited maternally from the ovaries through egg-smearing or through a yet-to-be-described form of transovarial transmission method [6,7]. However, this study did not characterize the bacterial communities associated with the ovarian tissues from either species to validate this possibility. Additional future studies characterizing the bacterial communities of mosquito ovaries in addition to the egg rafts and midgut tissues would shed more light on this question.

The midgut samples of *Cx. pipiens* and *Cx. restuans* mostly shared similar bacterial community composition with over 61% of the bacterial OTUs shared between them. Previous studies with the adults of these two mosquito species have generated variable results, with one study showing that they harbor distinctly different bacterial communities [47], while another study did not report unique clustering in the bacterial communities of the two species [48]. Both studies were conducted with non-blood-fed adult mosquito samples whose parity status was not assessed, a factor that might have contributed to the observed differences. We can partly attribute similarity in the bacterial OTUs for adults of the two species in this study to the shared environment resulting in colonization by similar bacterial communities. Mosquitoes from different species sampled from common habitats have been shown to share more similar bacterial communities, a possible indication of horizontal acquisition from the surrounding environment [8,48,49]. Similarly, the blood-fed and gravid status of the females at the time of sampling may have contributed to similar internal gut physiology subsequently supporting comparable bacterial communities [50,51].

The significant separation of the bacterial community composition between egg rafts and midguts within each mosquito species, is not surprising since the physiological environment in the mosquito gut is expected to differ substantially from that of the egg rafts, and thus is likely to facilitate colonization by very different consortia of bacterial communities. The disproportionate dominance of a few distinct bacterial taxa in the egg rafts compared to midgut samples for both species further indicates that egg and midgut environments were substantially different. However, studies comparing the bacterial communities of mosquito midguts and eggs are rare. One such study focusing on *Aedes aegypti* shows that mosquito guts share a significant proportion of their bacterial community composition with those of the eggs. Most of the taxa that have been reported to be shared between mosquito egg and midgut life stage are widespread in mosquito species and have been described in many other mosquito microbial studies [5].

There was high bacterial richness in the egg raft samples compared to midgut samples for both *Cx. pipiens* and *Cx. restuans*. The midgut samples were all from gravid females that are likely to have experienced a sudden and sharp decline in bacterial diversity due to their prior bloodmeal diet [50,52]. In mosquito vectors, bloodmeal diet is associated with significant reductions in the bacterial diversity of the mosquito gut. This is attributed to the breakdown of the heme proteins associated with the bloodmeal diet releasing reactive oxygen species that alter the physiological configuration of the gut environment allowing only bacterial taxa that can tolerate the high oxidative stress [50,52]. The digestive process involving the movement of the blood bolus along the midgut endoperitrophic space may also contribute to physical propulsion and excretion of a significant proportion of the midgut bacterial communities further reducing the midgut bacterial diversity.

The dominance of *Ralstonia* and *Novosphingobium* in egg raft samples of both species point to their possible adaptation to colonizing the mosquito egg raft stages. It also could be related to their potential role in mosquito ecology, such as inducing egg hatch, but this requires further research. Literature is scanty on the isolation and characterization of *Ralstonia* or *Novosphingobium* from mosquito egg stages, but this may be attributed to the dearth of studies characterizing mosquito egg bacterial communities. However, *Ralstonia* and *Novosphingobium* have been isolated in mosquito midgut samples from several mosquito species including *Aedes japonicus, Ae. aegypti, Ae. albopictus*, and *Anopheles coluzzii*. They have also been described from the natural environment, including soil and aquatic sources, indicating that mosquitoes may acquire them horizontally [53–58]. Their isolation from egg rafts in this study shows that these taxa may be part of the common mosquito bacterial commensals shared between different mosquito life stages including the eggs. Additional studies characterizing the bacterial communities of *Culex* ovaries, in addition to egg rafts, would help shed light on microbial presence in the ovary tissues and whether they are passed maternally to the egg rafts through egg-smearing or other forms of transovarial transmission.

The midgut samples of *Cx. pipiens* and *Cx. restuans* were dominated by *Providencia*, while *Wolbachia* and *Spironema* were the second most abundant bacterial taxa in *Cx. pipiens* and *Cx. restuans* midgut samples, respectively. We attribute the dominance of *Providencia* in both species potentially to the bloodmeal diet, an indication that this taxon could tolerate the high oxidative stress and the enzymatic conditions in the midgut environment. The genus *Providencia* has been isolated from several mosquito species, including *Anopheles albumanus* [59], *Aedes aegypti* [54,60], and *Aedes vexans* [48]. However, it has not been isolated in *Cx. pipiens* or *Cx. restuans* from previous bacterial studies. *Providencia* is a genus consisting of gram-negative rods with peritrichous flagella belonging to the family Enterobacteriaceae. It is an enteric bacterial pathogen commonly isolated from human intestines. It has also been isolated from other organisms, including birds and pigs, pointing to its ubiquity in the natural environment. However, its role in mosquito biology has not been described. The ease of culturing *Providencia* in bacteriological media, its ready availability in the natural environment, and its abundance in *Cx. pipiens* in this study points to its potential suitability as a candidate for manipulation for paratransgenesis for mosquito vector management [61]. The bacterial genus *Spironema* has been characterized in *Culiseta melanura* [62], *Ae. aegypti* un-infected with *Wolbachia* [63], *Cx. pipiens* [64], and was also dominant in *Culex nigripalpus* [65]. This taxon has been isolated from soil as well as river water samples, providing evidence of its potential horizontal acquisition in the *Cx. restuans* in this study [66]. High abundance in *Cx. restuans* has not been reported previously and opens an avenue to conduct further studies on its potential role in mosquito biology and disease transmission. The presence of *Wolbachia* in *Cx. pipiens* was expected as *Cx. pipiens* naturally harbor *Wolbachia*, a maternally inherited endosymbiont common in many arthropods, where they mediate several reproductive manipulations in their hosts [67]. Our study did not report any trace of *Wolbachia* in *Cx. restuans* egg raft samples. However, it was reported in the *Cx. restuans* midgut samples at <0.1%, providing further evidence of its recent detection in *Cx restuans* elsewhere [68]. The presence of *Wolbachia* in *Cx. restuans* and *Cx. pipiens* has potential to alter the epidemiology of WNV infection especially in the regions of the US where WNV has become endemic. Transient infection of *Cx. tarsalis* with the *w*AlbB strain has been associated with enhanced replication of WNV in the host [69]. In other studies, transient infection of *Ae. aegypti* with WNV was associated with enhanced viral replication [14]. However, it is likely that the interaction between *Wolbachia* and WNV in the naturally infected *Culex* populations could manifest differently. This area requires further investigation to disentangle the role of *Wolbachia* in WNV incubation and subsequently the epidemiology of the disease in the endemic regions.

In conclusion, our study has shown that *Cx. pipiens* and *Cx. restuans* egg raft samples are more diverse in their bacterial communities and the bacterial communities differ significantly between egg raft and midgut tissues within each mosquito species. However, the bacterial communities of the egg raft versus the midgut tissues of the two species are mostly similar in their community composition. Whereas previous studies with *Cx. pipiens* or *Cx. restuans* have prioritized characterizing the bacterial communities from adult midguts [47,48,70], the additional characterization of the egg raft bacterial communities in this study fills an important gap in our understanding of the bacterial communities associated with the egg rafts and how they compare with those of the midguts. These findings open the way for further studies on their role in mosquito biology and ecology and their potential to be exploited for mosquito vector management such as strategies to interfere with egg hatching in the environment.

## Declarations

### Availability of data and materials

The datasets used and/or analysed during the current study are available from the corresponding author on reasonable request.

### Competing Interests

The authors declare no competing financial or non-financial interests

## Funding

This research was supported by the U.S. Department of Agriculture, Agricultural Research Service. This work was also supported by NSF DEB 1754115; Waste Tire Fund and Emergency Public Health Act from the State of Illinois; University of Illinois Institute for Sustainability, Energy, and Environment grant. Any opinions, findings, conclusions, or recommendations expressed in this publication are those of the author(s) and do not necessarily reflect the view of the U.S. Department of Agriculture. Mention of trade names or commercial products in this publication is solely for the purpose of providing specific information and does not imply recommendation or endorsement by the U.S. Department of Agriculture. USDA is an equal opportunity provider and employer.

## Authors’ contributions

E.O.J, B.F.A, and C.S. conceived the study, E.O.J. conducted the experiment and analyzed the data, C.H.K assisted with conducting the experiments, C.D. processed the sequencing data and generated the OTU table, C.S. and C.D. contributed, reagents, materials, and analysis tools. All authors contributed to writing and approved the manuscript.

## Acknowledgments

We thank Juma Muturi, and Millon Blackshear for their technical assistance and all members of the Medical Entomology Laboratory (INHS), Yani Kaldis, Rosemary Ogbonna, and Eddie Lewis, for assisting with fieldwork and running the experiments, and members of Allan Laboratory for providing valuable reviews to the manuscript. The authors would like to thank Heather Walker of the National Center for Agricultural Utilization Research (U.S. Department of Agriculture) for helpful technical assistance. Any opinions, findings, conclusions, or recommendations expressed in this publication are those of the author(s) and do not necessarily reflect the view of the U.S. Department of Agriculture. The mention of firm names or trade products does not imply that they are endorsed or recommended by the USDA over other firms or similar products not mentioned. USDA is an equal opportunity provider and employer.

## Notes

### Competing Interest Statement

The authors have declared no competing interest.

