## Supplementary material for "*Culex pipiens* L. and *Culex restuans* egg rafts harbor diverse bacterial communities compared to their midgut tissues"

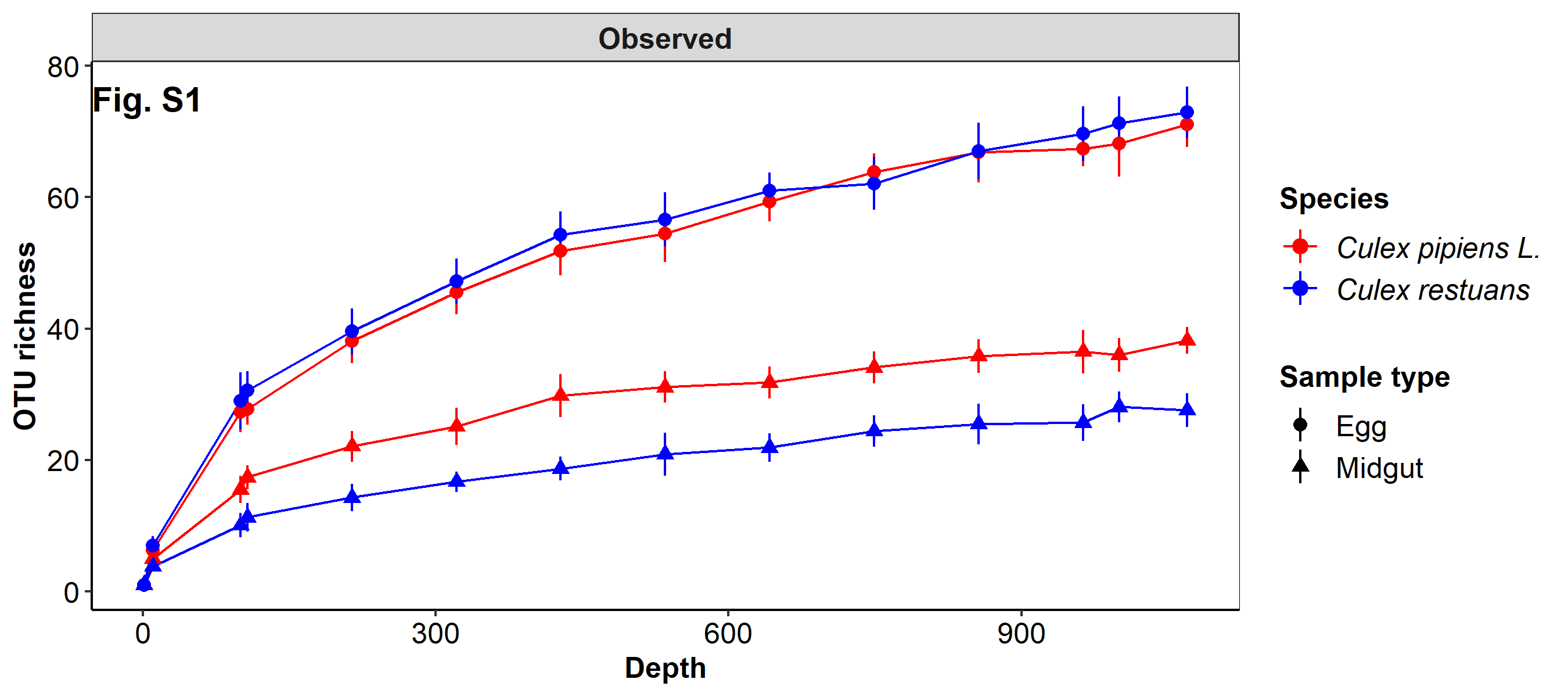

**Additional file 1: Fig. S1** Rarefaction curve analysis of observed richness of bacterial OTUs of samples from Cx. pipiens and Cx. restuans midgut and egg samples

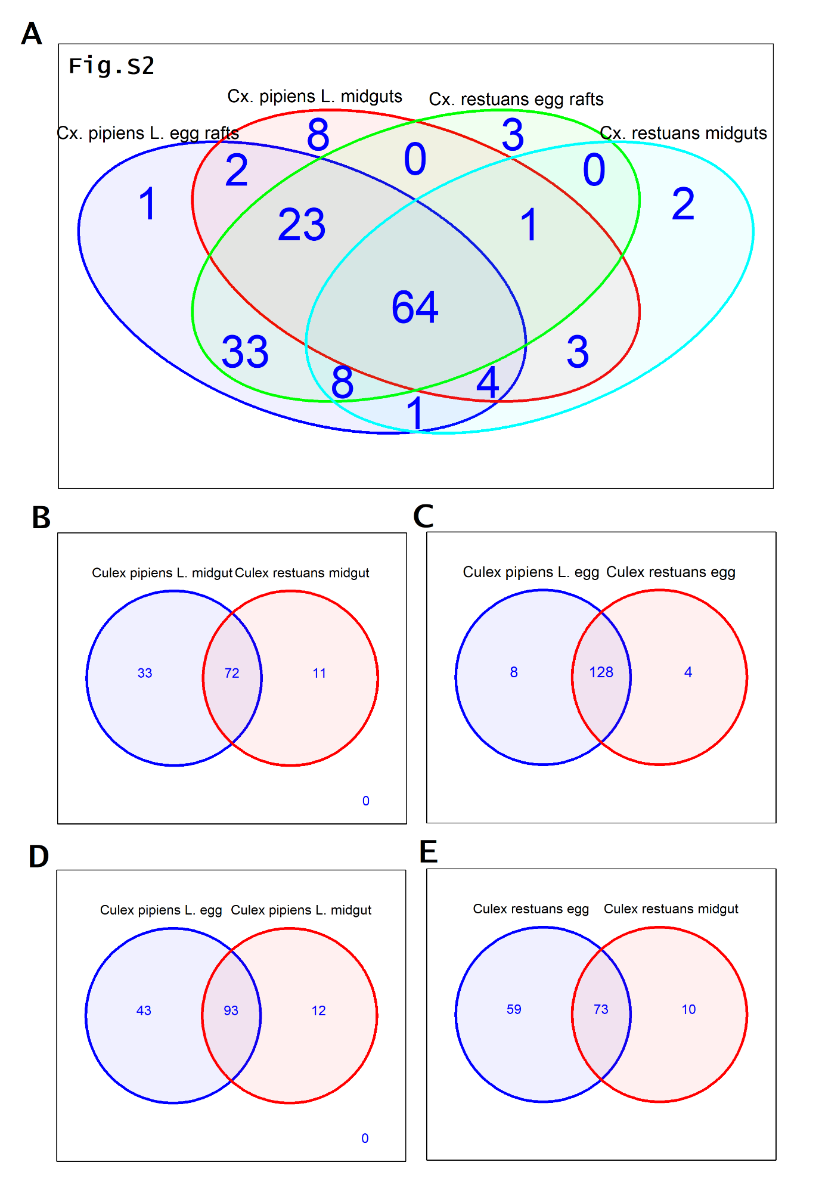

**Additional file 2: Fig. S2** Venn diagrams showing the number of unique and shared bacterial OTUs between egg and midgut samples of Cx. restuans and Cx. pipiens. A: venn analysis of bacterial OTUs from all four sample types; B: venn analysis of bacterial OTUs from Cx. pipiens and Cx. restuans midgut samples; C: venn analysis of bacterial OTUs from Cx. pipiens and Cx. restuans egg samples; D: venn analysis of bacterial OTUs from Cx. pipiens eggs and midguts; E: venn analysis of Cx. restuans egg and midgut samples

**Additional file 3: Table S1**. SIMPER analysis of the major bacterial OTUs driving differences between sample treatments (mosquito species and life stage). The Bray-Curtis average dissimilarity between sample treatments was >1% for 18 bacterial taxa. Overall average dissimilarity. CXP.EG – Cx. pipiens egg raft samples; CXP.MG – Cx. pipiens midgut samples; CXR.EG – Cx. restuans egg raft samples; CXR.MG – Cx. restuans midgut samples.

| **Taxon** | **Average dissimilarity** | **Percent contribution** | **Cumulative percentage** | **CXP.EG** | **CXP.MG** | **CXR.EG** | **CXR.MG** |
| --- | --- | --- | --- | --- | --- | --- | --- |
| *_Providencia_* | _17.11_ | _19.14_ | _19.14_ | _4_ | _4630_ | _7_ | _6060_ |
| *_Ralstonia_* | _9.453_ | _10.57_ | _29.71_ | _2010_ | _125_ | _1490_ | _47_ |
| *_Novosphingobium_* | _9.247_ | _10.34_ | _40.06_ | _1250_ | _900_ | _1880_ | _153_ |
| *_Spironema_* | _7.497_ | _8.386_ | _48.44_ | _6_ | _217_ | _79_ | _7000_ |
| *_Wolbachia_* | _7.454_ | _8.338_ | _56.78_ | _415_ | _2430_ | _0_ | _7_ |
| *_Acinetobacter_* | _4.563_ | _5.104_ | _61.88_ | _683_ | _118_ | _624_ | _565_ |
| _Methylobacteriaceae_ | _2.957_ | _3.307_ | _65.19_ | _163_ | _23_ | _1040_ | _19_ |
| *_Methylobacterium mesophilicum_* | _2.58_ | _2.886_ | _68.08_ | _478_ | _19_ | _371_ | _1_ |
| *_Flectobacillus_* | _2.503_ | _2.799_ | _70.88_ | _288_ | _152_ | _533_ | _9_ |
| _Enterobacteriaceae_ | _2.319_ | _2.593_ | _73.47_ | _28_ | _460_ | _174_ | _274_ |
| *_Erwinia_* | _2.163_ | _2.42_ | _75.89_ | _0_ | _109_ | _0_ | _871_ |
| *_Ramlibacter_* | _2.084_ | _2.331_ | _78.22_ | _293_ | _1_ | _399_ | _1_ |
| _Comamonadaceae_ | _1.551_ | _1.735_ | _79.95_ | _283_ | _49_ | _247_ | _3_ |
| *_Aquitalea_* | _1.337_ | _1.496_ | _81.45_ | _184_ | _40_ | _285_ | _0_ |
| *_Sphingomonas yabuuchiae_* | _1.22_ | _1.365_ | _82.81_ | _239_ | _3_ | _136_ | _2_ |
| *_Serratia_* | _1.075_ | _1.202_ | _84.02_ | _39_ | _218_ | _0_ | _24_ |
| *_Novosphingobium capsulatum_* | _1.031_ | _1.153_ | _85.17_ | _176_ | _28_ | _191_ | _8_ |
| _Methylophilaceae_ | _1.016_ | _1.137_ | _86.31_ | _170_ | _0_ | _167_ | _1_ |
